# Cholesterol modulates the human FPN1 iron export function in plasma membrane liquid-ordered microdomains

**DOI:** 10.1101/2023.12.14.571614

**Authors:** Rim Debbiche, Chandran Ka, Isabelle Gourlaouen, Sandrine Maestri, Kévin Uguen, Paul-Alain Jaffrès, Isabelle Callebaut, Gérald Le Gac

**Author notes:** **Corresponding Author:** Gerald Le Gac, Institut Brestois de Recherche en Bio-Santé, UFR Médecine et Sciences de la Santé, UMR1078, 22 rue Camille Desmoulins, 29238 Brest, France. **Authorship contributions** G. Le Gac, I. Callebaut and PA. Jaffrès designed the study. R. Debbiche, C. Ka, I. Gourlaouen, S. Maestri, K. Uguen and I. Callebaut conducted experiments. R. Debbiche, C. Ka, I. Callebaut, PA. Jaffrès and G. Le Gac analyzed data. G. Le Gac and I. Callebaut wrote the manuscript. All authors contributed to the editing of the final manuscript.

## Abstract

Ferroportin 1 (FPN1) is the only known mammalian iron efflux transporter. This multi-pass membrane protein, which adopts the Major Facilitator Superfamily fold, is tightly controlled by serum hepcidin to assure maintenance of adequate cellular and systemic iron levels. Earlier studies have shown that cholesterol-lowering drugs can reduce FPN1 expression in liquid-ordered plasma membrane microdomains and its sensitivity to hepcidin. However, the molecular mechanism by which cholesterol depletion regulates the localization of FPN1 at the cell surface remains unknown. In biochemical experiments, we show that cholesterol depletion reduces the iron export function of FPN1. Repletion with cholesterol restores FPN1 activity. This is not observed with the diastereoisomer epicholesterol, suggesting a direct interaction between cholesterol and FPN1. Consistent with this, we demonstrate that mutants affecting the key tyrosine residues of three cholesterol-recognition amino acid consensus (CRAC/CARC) motifs have a negative impact on FPN1 activity, in manner that also decreases its abundance in ordered plasma membrane microdomains. A complementary structural analysis allows us to focus on a conserved CARC motif (CARC-1) located in a deep hydrophobic groove between transmembrane helices 1 and 5. Molecular docking suggests that this groove is well suited to cholesterol binding. All these findings indicate that the interaction between FPN1 and cholesterol is of major importance for the localization of FPN1 in ordered microdomains of the plasma membrane, which is necessary for its optimal activity, and so its responsiveness to hepcidin.

## INTRODUCTION

Mammalian cells utilize several mechanisms to acquire iron, but ferroportin 1 (FPN1 - also known as solute carrier family 40 member 1/SCL40A1 - ; Uniprot ID: Q9NP59) is the only way out (1, 2). FPN1 is therefore essential for iron homeostasis, at both cellular and systemic levels (3). Although many different regulatory mechanisms have been documented at the DNA, RNA and protein levels, FPN1 is predominantly regulated by the liver-derived peptide hepcidin which, depending on the cell type, induces internalization and degradation of FPN1 and/or blocks the FPN1-related iron export mechanism (4, 5). The FPN1-hepcidin axis is associated with inherited and acquired disorders of iron metabolism (6, 7), while FPN1 down-regulation has been reported in various cancer cells, where it is believed to promote iron retention and cell proliferation (8).

FPN1 belongs to the Major Facilitator Superfamily (MFS), a large group of secondary active transporters that is present in cells across all life kingdoms, controlling the flow of a broad range of substrates (inorganic ions, metabolites, neurotransmitters, and drugs) over lipid bilayers. MFS proteins share a common architecture consisting of two bundles of six transmembrane (TM) helices (N-domain: TM1-TM6; C-domain: TM7-TM12), related by twofold pseudosymmetry (9–12). Since the seminal paper published by Jardestzky et al. in 1966 (13), many researchers have sought to elucidate the molecular basis of conformational changes in MFS proteins in different cellular environments and species. Nowadays, there seems to be a general consensus in favor of the “clamp-and-switch” transport cycle model (18, 19). A better understanding of the role of lipids in the fine-tuning of MFS proteins is now identified as a high priority goal by some authors, who have also highlighted some important technical issues, particularly for mammalian proteins (14).

One of the most striking features of FPN1 is a preferential localization in low-density fractions of detergent-resistant plasma membrane microdomains (DRM) of cells that handle large amounts of iron (15–17). This particular environment, which corresponds to liquid-ordered states of biological membranes enriched in cholesterol and sphingolipids and is referred to as lipid-rafts by many authors (18–23), has also been proven critical for endocytosis and degradation of FPN1 after hepcidin binding in mouse macrophages. Interestingly, hepcidin activity was diminished after treatment with filipin and methyl-beta-cyclodextrin, two different compounds that cause liquid-ordered phases breakdown by cholesterol depletion (15).

It is important to emphasize that cholesterol (Chol) is a major constituent of eukaryotic plasma membranes that has a major impact on membrane organization, fluidity and dynamics. Chol can promote the formation of liquid-ordered microdomains (which have a higher Chol concentration), where it is thought to modulate the structure and function of some embedded proteins through direct interaction with them and/or changes in the biophysical properties of the plasma membrane (20, 22, 24–29). Typical Chol recognition amino acid consensus (CRAC) motifs, which contain amino acids that can interact with Chol by hydrogen bonds (hydroxyl group of Chol vs. Arg or Lys) and hydrophobic (alkyl group of Chol vs. Leu or Val) and stacking interactions (sterane backbone of Chol vs. Tyr, Phe or Trp) (30), have been identified in mouse and human FPN1, suggesting a strong potential for direct binding of Chol (15). However, whether the iron export function of FPN1 depends on interactions with surrounding Chol molecules in the lipid bilayer is yet to be established.

In this study, we show that human FPN1 is modulated by Chol in liquid-ordered microdomains of the plasma membrane of human embryonic kidney cells. Sequence analysis indicate that FPN1 contains three major Chol-binding motifs, which are supported by site-directed mutagenesis of aromatic amino acids and biochemical assays. The three considered motifs are critical in mediating the abundance of FPN1 in Chol-enriched microdomains, in a manner consistent with full iron export capacity.

## METHODS

### FPN1 plasmid constructs

The wild-type (WT) FPN1-V5 and FPN1-V5/CD8 bicistronic plasmid constructs were generated as previously described (31, 32). The full-length human FPN1 cDNA (Genbank accession number NM_0414585.5) was inserted into the pcDNA5-FRT-TO vector (ThemoFisher Scientific) to produce isogenic stably transfected tetracycline-inducible human epithelial kidney 293 cells (Flp-In 293 T-Rex). All FPN1 mutations were introduced in the different vectors by using the QuikChange Site-Directed mutagenesis kit, according to the manufacturer’s instructions (Agilent Technologies). Sequencing analyses were performed to check the integrity of all plasmid constructs (full length SLC40A1 cDNA sequenced after each site-directed mutagenesis).

### Culture and transfection of human epithelial kidney (HEK)293T cells

Human epithelial kidney (HEK) 293T cells, from the American Type Culture Collection, were incubated at 37°C in a 5% CO_2_ humidified atmosphere and propagated in Dulbecco’s modified Eagle’s medium (DMEM; Lonza) supplemented with 10% fetal bovine serum. Cells were transiently transfected using Lipofectamine 2000 (ThermoFisher Scientific; flow cytometry experiments) or Transit-2020 (Mirus Bio LLC; DRM and iron export experiments), according to the manufacturer’s instructions, and a 3:1 transfection reagent (μL)/plasmid DNA ratio (μg).

### Generation of stable HEK293 cells exhibiting tetracycline-inducible expression of human FPN1

Flp-In 293 T-Rex cells were transfected with the different pcDNA5-FRT-TO plasmid constructs and the pOG44 vector using the TransIT2020 cationic lipid (Mirus Bio LLC), according to the manufacturer’s instructions (ThemoFisher Scientific). Stably transfected cells were selected in DMEM (Lonza) supplemented with 10% fetal bovine serum, 150 μg/mL blasticidin (ThermoFisher Scientific) and 300 μg/mL hygromycin (ThermoFisher Scientific) for 3 weeks. The resulting clones were propagated and maintained in the same media. *SLC40A1* cDNA integration was verified by PCR amplification and Sanger sequencing, while FPN1 expression was assessed by Western-Blotting.

### Isolation of detergent resistant membranes

Transiently transfected HEK293T or stably transfected HEK293 cells were washed ice-cold phosphate buffered saline (PBS), scraped into 2 mM EDTA/PBS, centrifuged (200 x g, 5 min, 4°C), and incubated on ice for 60 min in 1 mL lysis buffer (10 mM Tris HCl pH7, 10 mM EDTA, 100 mM NaCl; 1% Triton X-100) supplemented with protease inhibitor cocktail (Roche) and phenylmethanesulfonyl fluoride (PMSF; Sigma Aldrich). Cell lysates were homogenized by 20 passages through a 25-gauge needle and centrifuged at 1,200 x g for 10 min at 4°C. The supernatants were adjusted to a final concentration of 40% (w/v) iodixanol (OptiPrep, Sigma Aldrich), and the mixtures were then layered under a 20-40% iodixanol discontinuous gradient before to be centrifuged at 260,000 x g for 16h at 4°C using an SW41 Ti rotor (Beckman Coulter). After spinning, eleven fractions of 1 mL were collected from the top to the bottom of the gradient tube, and Western blot analysis of the fractions was performed. Detergent-Resistant Membrane (DRM) and Detergent-Sensitive Membrane (DRS) fractions were defined by the presence of flotillin 1 and TfR1 markers (15), respectively.

### Western blotting analysis

Collected fractions from iodixanol gradient (25 μL) were solubilized in Laemmli buffer, resolved by a sodium dodecyl sulfate-polyacrylamide gel electrophoresis (SDS-PAGE) and electro-transferred on 0.45 μM nitrocellulose membranes (GE Healthcare). The membranes were incubated in blocking solution (5% skim milk in 0.05% Tween-20/Tris Buffered Saline; TBST) for 1h at room temperature, and incubated overnight at 4°C with primary antibodies: rabbit anti-V5 (Cell signaling, D3H8Q; 1/1000); rabbit anti-flotillin 1 (Cell signaling, D2V7J; 1/1000), rabbit anti-TfR1 (Cell signaling, D7G9X; 1/1000). After three washes with TBST, blots were incubated for 1h at room temperature with a horse radish peroxidase-labeled secondary antibody (anti-rabbit, Dako; 1/1000), and revealed by ECL chemiluminescence (Millipore) and in a GeneGnome digital imaging system (Syngene).

### Plasma membrane cholesterol modulation in human SLC40A1 stably transfected HEK293 cells

Methyl-β-cyclodextrin (MβCD; Sigma) was dissolved in a serum free media (Pro293A-CDM, Lonza), whereas sterols (Steraloids) were dissolved in a mixture of chloroform-methanol (1:1, v/v). MβCD:sterol complexes were prepared by adding 0.1 mM of cholesterol or epicholesterol to 1 mM MβCD (1:10 dilution) (33), and repeatedly vortexing the mixtures for 1h at 37°C.

Plasma cholesterol was depleted by treating HEK293 cells with 5 mM MβCD for 30 min in Pro293A-CDM (Lonza); this short period of MβCD treatment was deemed sufficient to removed 50% of endogenous cholesterol from cells (34). Cholesterol and epicholesterol replenishments were performed by first depleting the membrane cholesterol with MβCD (5 mM, 30 min), followed by incubation of HEK293 cells with MβCD:cholesterol or MβCD:epicholesterol complexes (10:1 mM) for 3h. FPN1 expression was induced following these treatments. All experiments were carried out by maintaining the cells at 37°C in a humidified atmosphere with 5% CO_2_.

### ^55^Fe release measurements

^55^Fe loading of human apotransferrin was performed as previously described (35). Briefly, HEK293T cells (1.7 x 10^5^ cells/well in 12-well plates) were grown for 24 h in supplemented DMEM (Lonza), before to be incubated with 20 μg/mL ^55^Fe-transferrin for 24 h and transiently transfected with wild-type or mutated FPN1-V5 plasmid constructs. Fifteen hours post-transfection cells were washed once with PBS and cultured in Pro293a-CDM serum-free medium (BioWhittaker) for up to 15 (cholesterol depletion/sterols repletion experiments) or 36 h (characterization of FPN1 mutants). ^55^Fe exported into the supernatant was collected, mixed with liquid scintillation fluid (Ultima Gold MV, Packard Bioscience) and counted for 10min in a TRICARB 1600 CA scintillation counter (Packard). Percentage of ^55^Fe export was calculated using the following formula: (^55^Fe in the supernatant at end point, divided by cellular ^55^Fe at time zero) x 100.

### Flow cytometry analysis

Flow cytometry experiments were done as previously described (32). Briefly, HEK293T cells (1.75 x 10^5^ cells/well in 6-well plates) transfected with the pIRES_FPN1-V5_CD8 constructs were treated (or not) with 4.3 nM of native human-25 hepcidin (24h after transfection) for 16 hours. Cells were harvested with trypsin, pelleted (500 g, 5 min, 4°C) and resuspended in PBS (pH 7.4) containing EDTA (Lonza) and 10% fetal bovine serum, before being incubated for 20 min at 4°C with anti-V5-FITC or anti-V5-PE (ThermoFisher Scientific) and anti-CD8-APC (Miltenyi Biotec). Stained cells were pelleted (500 g, 5 min, 4°C) and resuspended in 400 *μ*L PBS-EDTA. Cells were analyzed using a BD Accuri C6 flow cytometer (BD Biosciences) and FlowLogic™ software (Miltenyi Biotec).

### 3D structure analysis

The experimental 3D structure of human ferroportin in the outward-facing state (6W4S; apo configuration) was extracted from the Protein Data Bank (https://rcsb.org/) and manipulated using Chimera 1.13.1. (36). Blind docking was performed using the EADock DSS tool provided by the online SwissDock server (37, 38). The 3D structure was positioned in a lipid bilayer using the PPM web server (39).

### Statistical analysis

Data are presented as means (column bars) + standard deviation. Comparison used 2-tailed paired sample *t*-test.

## RESULTS

### FPN1 predominantly localizes in detergent-resistant plasma membrane microdomains of HEK293T cells

Mouse Fpn1 has been reported to preferentially localize in detergent-resistant membrane domains (DRM) of the plasma membrane of bone marrow derived macrophages isolated from DBA2 mice. Fpn1 expression has even been shown to be increased in these domains after iron treatment (Fe-NTA) (15).

We aimed to reproduce these findings in HEK293T cells, transiently transfected to express a wild type human FPN1-V5 fusion protein. Flottilin-1, a marker of DRM fractions, was mainly detected in fractions 1-4, whereas TfR1, a marker of non-DRM fractions, was mainly detected in fractions 9-12. Flottilin-1 and TfR1 DRM profiles remained the same following treatment with 1 mg/mL holo-transferrin for 48h. The prevalent enrichment of human FPN1 in liquid-ordered fractions of the plasma membrane was confirmed, with more pronounced staining of the first four fractions after iron treatment (Figure 1). Similar results were obtained from T-Rex-293 cells, stably transfected to overexpress the human FPN1 protein (data not shown).

**Figure 1.**
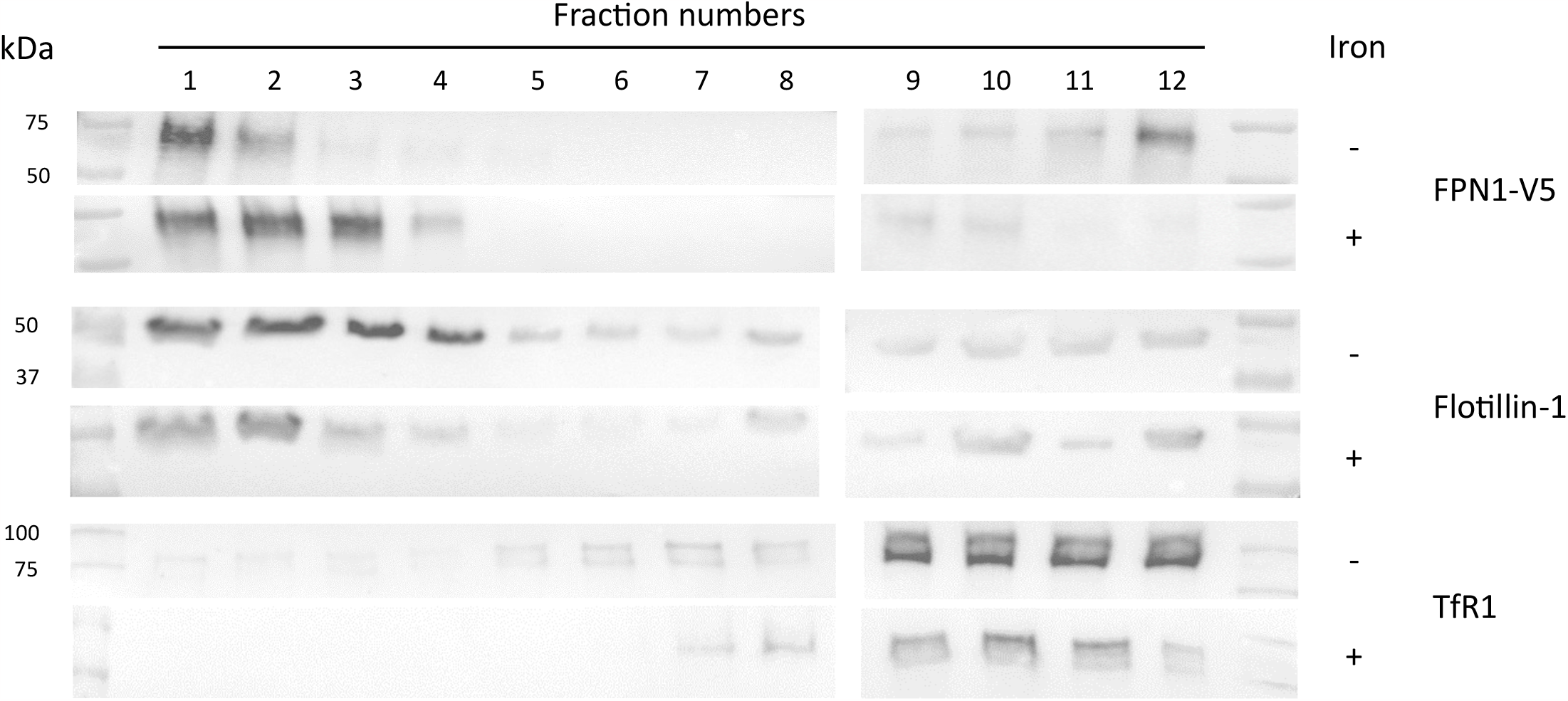
Distribution of human FPN1 in the plasma membrane of HEK293T cells treated or not with iron. Transiently HEK293T transfected cells were lysed in ice-cooled 1% Triton-X100 buffer. The mixture was loaded on a discontinuous iodixanol gradient and ultracentrifuged. Twelve 1mL fractions were collected from top (low density) to the bottom of the tube (fractions 1 to 12). Collected fractions were analyzed by Western blot using specific antibodies against V5-epitope (FPN1), Flotillin-1 (a marker of detergent-resistant microdomains) or TfR1 (a marker of detergent-sensitive microdomains). Flottilin-1 is mainly detected in fractions 1 to 5, where TfR1 is absent. FPN1 is enriched in detergent-resistant membrane microdomains after pre-treatment of HEK293T cells with holo-transferrin (1mg/mL holo-transferrin for 48h).

### FPN1 activity in the plasma membrane is dependent on cholesterol

Chol is a major component of liquid-ordered microdomains that can modulate the function of a wide range of proteins *via* specific and non-specific mechanisms. To determine whether FPN1 is responsive to surrounding Chol, we measured Fe^2+^ release from stably transfected T-Rex-293 cells expressing, upon induction by tetracycline, a wild-type FPN1-V5 fusion protein. The cells were incubated with MβCD or 10:1 MβCD:Chol to modify membrane Chol content.

As shown in Figure 2A, depletion of plasma membrane Chol resulted into a ∼30% reduction of FPN1 activity (p=0.0001). The iron export function was significantly and almost totally restored (∼6% reduction; p=0.055) by Chol repletion. Interestingly, such rescue of FPN1 activity was not observed with epicholesterol (∼26% reduction; p=0.003), which is the 3’ epimer of Chol (40). This inversion of chirality concerns the carbon atom carrying the hydroxyl group that is orientated on the α-face of the sterol instead of the β-face (Figure 2B). It is therefore not likely that a Chol-binding site in FPN1 also binds Epi-Chol.

**Figure 2.**
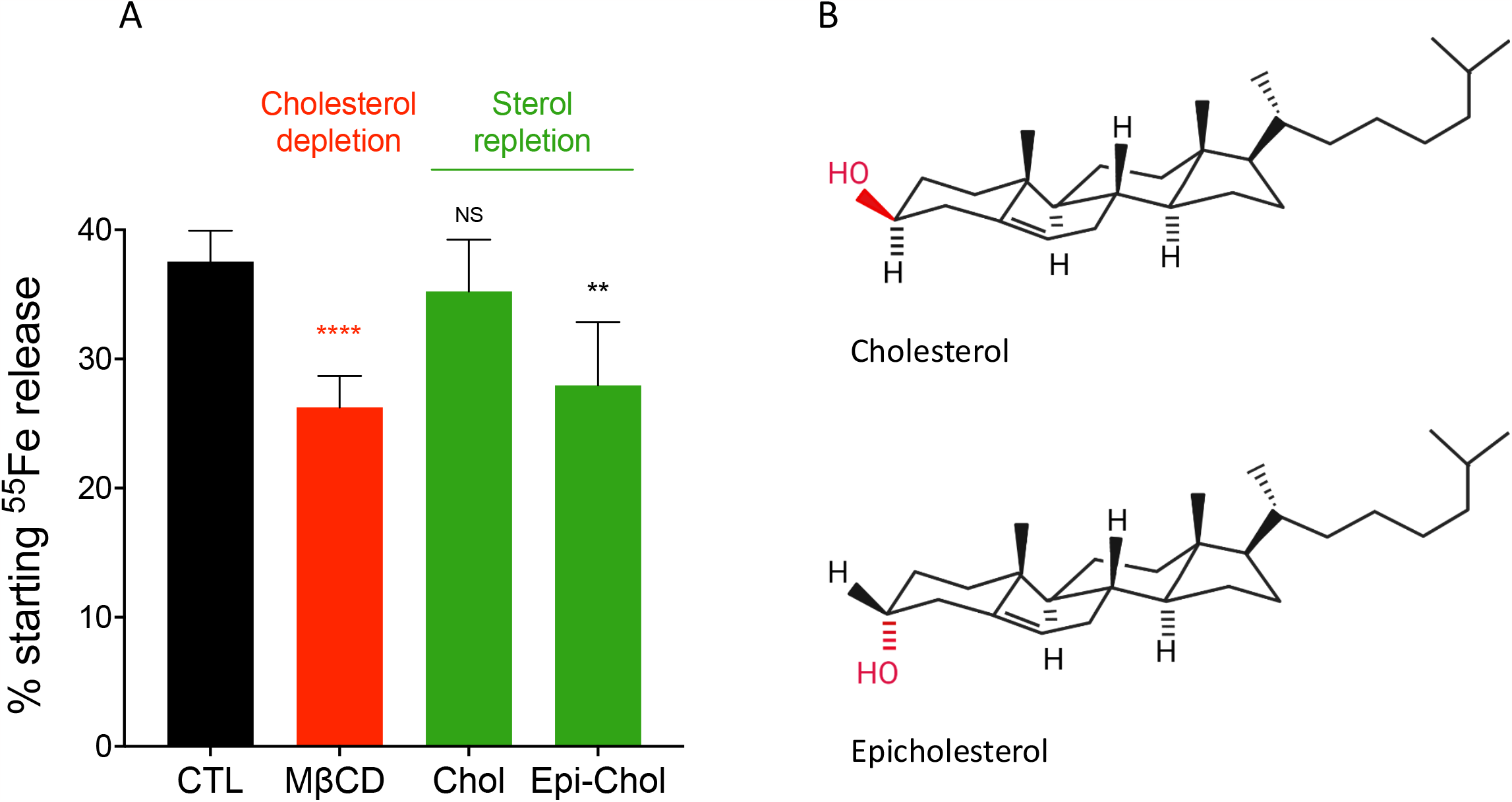
Membrane cholesterol alters FPN1 iron export function. (A) Flp-In™ T-Rex™ 293 cells were grown in 20 μg/mL ^55^Fe-transferrin for 24h before being washed and incubated or not (CTL) with MβCD (5 mM) for 30 min, and then with 10:1 MβCD:Cholesterol or MβCD:Epicholesterol for 3 h. FPN1 expression was induced after cholesterol depletion -/+ sterol repletion for 15 h (time zero), and the ^55^Fe exported into the supernatant was collected at 15 h post-induction. Data are presented as percentage of cellular radioactivity at time zero. Each bar represents the means ± sd of 5 independent experiments. P values were calculated by a Student’s t test. ** p<0.01 and **** p<0.0001. (B) Molecular structures of cholesterol and epicholesterol (α-3-OH-cholesterol epimeric form).

These results demonstrate that Chol modulates the plasma membrane function of FPN1, and further indicate that there is stereospecificity in the ability of Chol to promote iron export.

### FPN1 contains three putative cholesterol-binding motifs

To date, three major Chol-binding motifs have been identified in transmembrane proteins: one is conformational (Chol Consensus Motif; CCM), whereas the others two (CRAC and CARC) follow linear patterns. The CRAC motif (Chol Recognition/interaction Amino acid Consensus sequence), which is defined by the (L/V)-X_1-5_-(Y)-X_1-5_-(K/R) sequence from the N-terminus to the C-terminus, and its inverted and slightly modified version CRAC (N-ter-(K/R)-X_1-5_-(Y/F/W)-X_1-5_-(L/V) -C-Ter), have been the first to be described (and therefore also the ones that have been most extensively studied; (25)). In these motifs, positively charged basic amino acids (K/R) can form a hydrogen bond with the hydroxy group of Chol, which is supposed to be located at the lipid bilayer/water interface, whereas aromatic acids (Y/F/W) can bind to one of the four ring of the sterane backbone of Chol through CH-π staking, and hydrophobic amino acids (L/V) can form van der Waals interactions with the iso-octyl chain of cholesterol (25). In CCM, the amino acids that interact with Chol are distributed between two transmembrane helices (which interact with both sides of cholesterol): one helix contains the (W/Y)(I/V/L)(K/R) sequence, whereby the positively charged basic, aromatic and hydrophobic residues must face the same side of the helix (to bind Chol from one side), and one helix contains an aromatic residue (either phenylalanine or tyrosine; to bind Chol from the other side) (41, 42).

Focusing on the 12 transmembrane helices (TM) of the human FPN1 protein, which cross the lipid bilayer, and also taking into account their orientations with regard to Chol molecules within the inner and outer leaflets, we identified one CARC and two CRAC motifs. We named these motifs CARC-1 (TM1) and CRAC-1 and CRAC-2 (TM6) in Figure 3A. We also identified a putative CCM motif between transmembrane helices TM8 and TM10, which fits the features previously defined for the human α-adrenergic receptor (41) and the HA protein of the H7 influenza virus (42) (Supplementary Figure 1).

**Figure 3.**
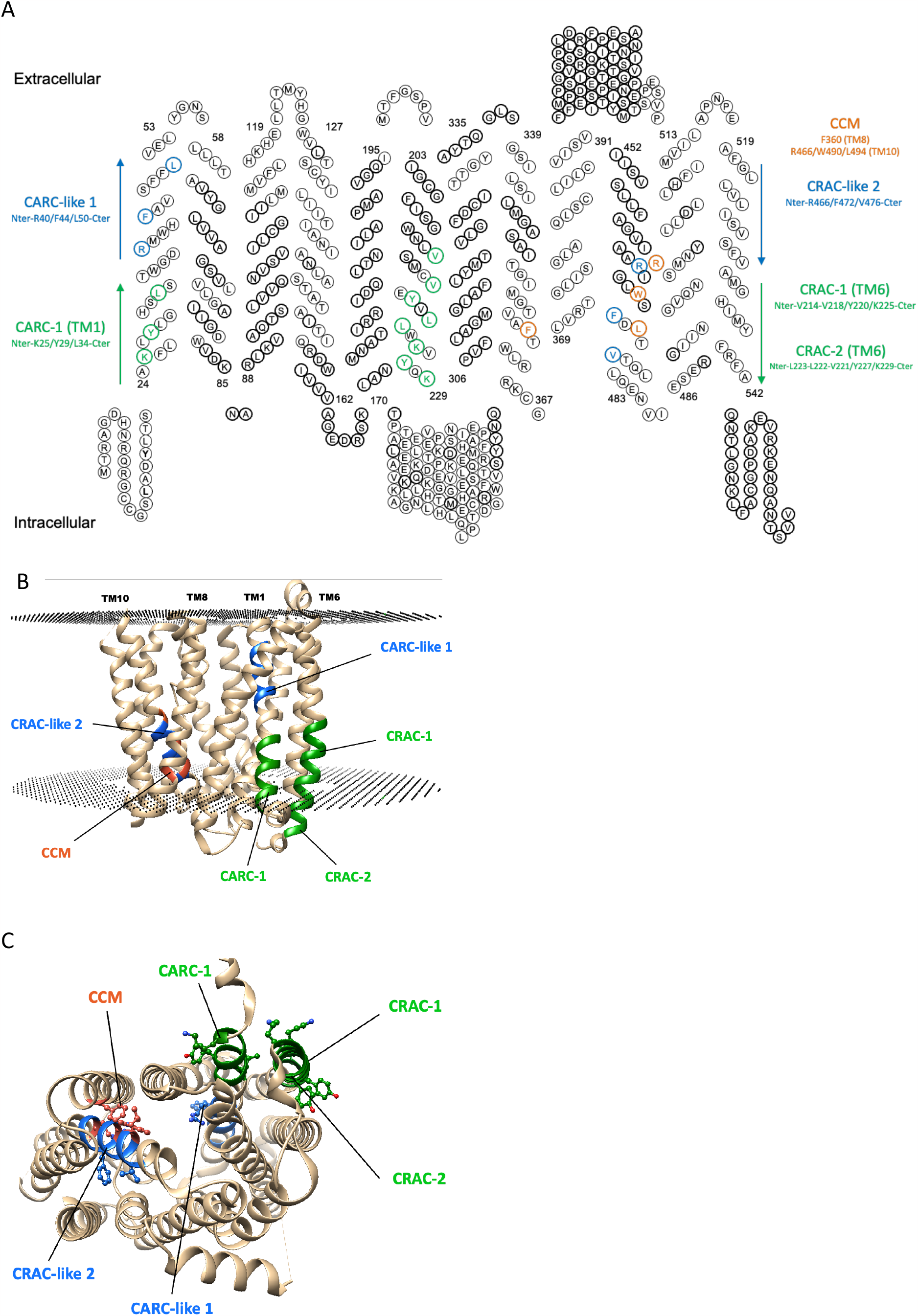
FPN1 contains potential CRAC, CARC and CCM cholesterol-binding motifs. (A) Two-dimensional representation of the human FPN1 amino acid sequence. Three putative Chol recognition motifs CRAC and CARC are present in transmembrane helices TM1 and TM6, respectively. The motifs are colored in green, and the characteristic residues K/R, Y and V/L are shown in bold enlarged circles. The CCM and CARC-like 1 and CARC-2 motifs are also represented and colored in orange and blue, respectively. (B-C) View of the 3D structure of human FPN1 in an outward-facing conformation (ribbon representation), in which the Chol recognition motifs are colored 5B), with the characteristic residues shown in atomic details (C). Two orthogonal views are displayed. The protein was positioned in a lipid bilayer using the PPM web server (39).

As shown in Figures 3B and 3C, the three linear Chol-binding sites of TM1 and TM6 are accessible to lipids at the surface of the FPN1 3D structure, but not the conformational CCM, which is buried within the TM assembly. This led us to consider only CARC-1, CRAC-1 and CRAC-2 as putative Chol binding sites.

### Specific mutants in predicted CARC and CRAC motifs reduce FPN1-mediated iron export

To verify the functional importance of the CARC-1, CRAC-1 and CRAC-2 motifs, we performed site-directed mutagenesis of the key central aromatic amino acids; the central tyrosine (Tyr29, 220 or 227) was substituted by either a phenylalanine (*i*.*e*. another aromatic amino acid) or a leucine (i.e. an aliphatic hydrophobic amino acid). Two unusual CARC motifs (not properly oriented, with positively charged basic amino acids buried in the hydrophobic core of the transmembrane helix) were identified in TM1 and TM10 (Figure 3A). They were named CARC-like 1 and CARC-like 2 and were also studied (Phe44 and Phe472 were changed for a leucine). The single amino acid Val162del deletion, which is known to reduce cell surface localization of FPN1 (35), was used as negative control in all experiments.

A bicistronic construct was used to evaluate the concurrent plasma membrane expression of the FPN1-V5 fusion proteins and cluster of differentiation 8 (CD8) in transiently transfected HEK293T cells on flow cytometry (Figure 4A). As expected, the level of FPN1-Val162del expression on the surface of co-transfected CD8+ cells was markedly decreased (∼25% reduction versus WT-FPN1; p<0.0001). More moderate reductions were observed with the Tyr220Phe, Tyr220Leu, Tyr227Leu mutants (<10% reduction; p<0.05), whereas no difference was observed with either the Tyr29Phe, Tyr29Leu, Tyr227Phe, Phe44Leu or Phe472Leu mutant.

**Figure 4.**
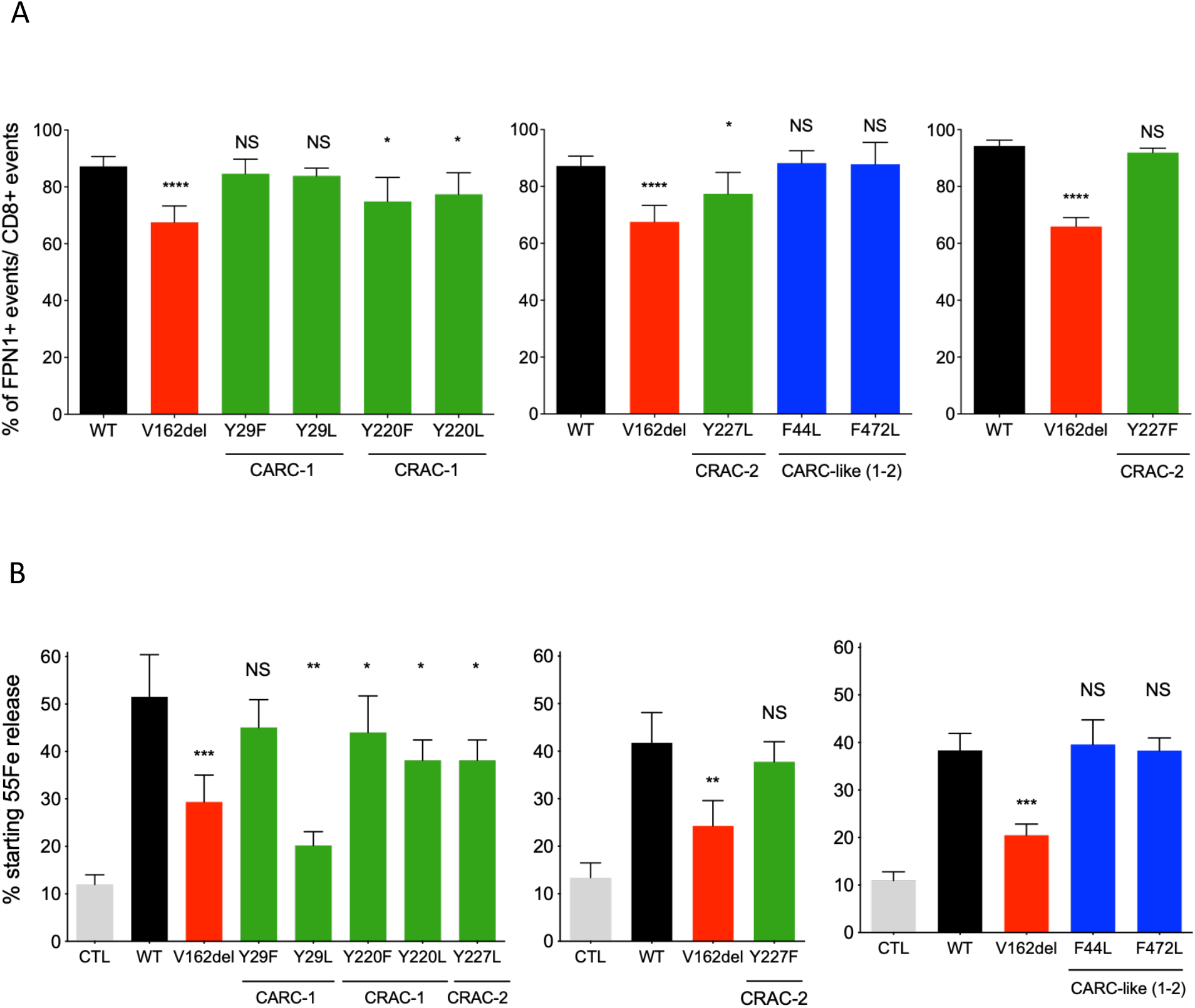
Mutational analysis of the cholesterol binding motifs CARC and CRAC in FPN1. (A) HEK293T cells were transiently transfected with the bicistronic pIRES2 plasmid encoding both full-length human FPN1-V5 and CD8. After 36 h, cells were double-stained for CD8 (APC) and the FPN1-V5 fusion protein (FITC, or PE) and analyzed by 2-color flow cytometry. Data are presented as percentages of FPN1-positive over CD8-positive events. (B) HEK293T cells were grown in 20 μg/mL ^55^Fe-transferrin for 24h before being washed and transiently transfected with wild-type or mutated SLC40A1-V5 expression plasmids. After 15 h, cells were washed and then serum-starved. The ^55^Fe exported into the supernatant was collected at 36h. Data are presented as percentage of cellular radioactivity at time zero. Each bar represents the means ± sd of 5 independent experiments. P values were calculated by a Student’s t test. ** p<0.01, *** p<0.001 and **** p<0.0001.

The activity of FPN1 mutants was then assessed in experiments measuring the export of radioactively labeled iron into the supernatant of transiently transfected HEK293T cells over a 36h period. As shown in Figure 4B, cells transfected with a pcDNA3.1 vector encoding a WT FPN1-V5 fusion protein showed 3-4-fold higher iron release than cells transfected with the corresponding empty vector (commercial pcDNA3.1-V5-His; No-FPN1). The Tyr220Phe, Tyr220Leu, Tyr227Leu mutants were not able to export ^55^Fe in amounts comparable with WT FPN1 (p<0.05), but were much more active than the Tyr29Leu mutant, which displayed activity similar to that of the well-known Val162del mutant (p<0.01).

These mutation data suggest that CARC-1, CRAC-1 and CRAC-2 motifs are important structural requirements for membrane Chol binding to FPN1, with larger effects for the changes of tyrosine to leucine, particularly at the Tyr29 position (CARC-1).

### The Tyr29Leu, Tyr220Leu, and Tyr227Leu mutants delocalize FPN1 from the liquid-ordered detergent-resistant microdomains of iron loaded HEK293T cells

As we did not observe any major effects of the tested mutants on the cell surface expression of FPN1, we finally asked whether the loss of the aromatic amino acid tyrosine at the central position of the CARC-1, CRAC-1 and CRAC-2 motifs of the TM1 and TM6 transmembrane helices has a consequence on the preferential localization of the iron exporter into detergent-resistant membrane microdomains.

To enhance FPN1 activity at the plasma membrane, the transiently transfected HEK293T cells were preloaded with 1 mg/mL holo-transferrin for 48h. Under these basal conditions, we confirmed the results presented in Figure 1 for the wild type protein (preferential localization in fractions 1-4). Interestingly, the Tyr29Leu mutant was found to totally shift to detergent-sensitive microdomains of the plasma membrane (very similarly to the TfR1 maker), which are known to contain lower concentrations of cholesterol (Figure 5). This effect was also apparent for the Tyr220 and Tyr227 mutants, but to a lesser extent and in a fairly gradual manner.

**Figure 5.**
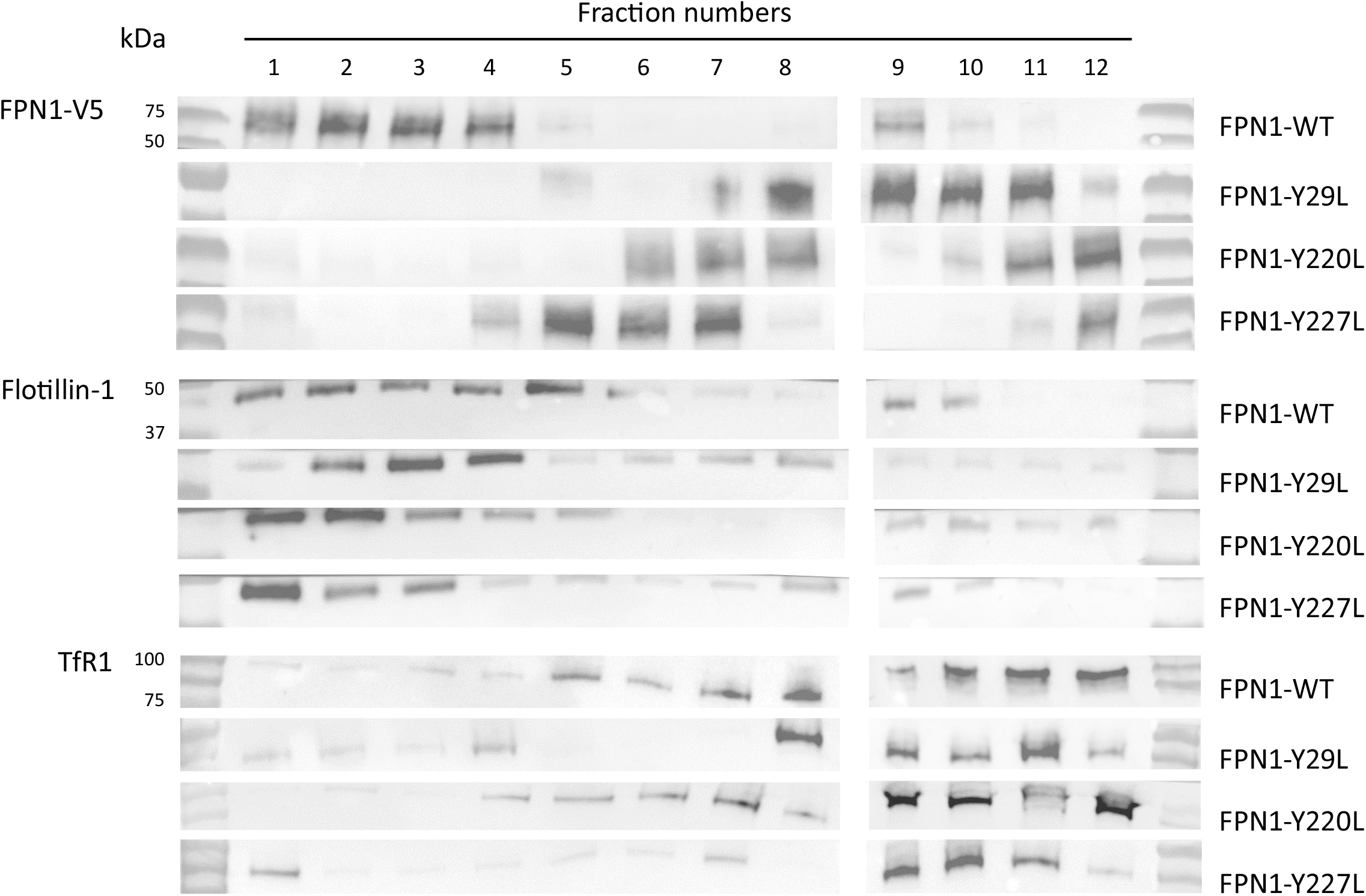
The Tyr29Leu, Tyr220Leu and Tyr227Leu mutants delocalize FPN1 from detergent-resistant membrane microdomains. Transiently HEK293T transfected cells were grown in 1mg/mL holo-transferrin before and then lysed in ice-cooled 1% Triton-X100 buffer. The mixture was loaded on a discontinuous iodixanol gradient and ultracentrifuged. Twelve 1mL fractions were collected from top (low density) to the bottom of the tube (fractions 1 to 12). Collected fractions were analyzed by Western blot using specific antibodies against V5-epitope (FPN1), Flotillin-1 (a marker of detergent-resistant microdomains) or TfR1 (a marker of detergent-sensitive microdomains).

Thus, the incorporation of the CARC and CRAC mutants into Chol-enriched microdomains of the plasma membrane is markedly decreased. This very likely explain the reduced capacity to export iron.

## DISCUSSION

Chol-enriched plasma membrane microdomains are thought to have specific functions in cell signaling, membrane transport, cell metabolism and migration, as well as pathogen invasion and disease (20, 22, 24–29, 43). Integral membrane proteins can be influenced by cholesterol in two non-exclusive ways: direct, based on an interaction between Chol and one or more Chol-binding sites, and indirect, based on various properties (fluidity, rigidity, thickness, lateral pressure, lipid order, dipole potential) of the membrane (44). Here, we identify tyrosine residues in candidate CARC and CRAC motifs as key determinants of the cell surface localization and iron export activity of human FPN1.

Auriac et al. previously reported that the iron exporter localizes to non-ionic detergent-resistant microdomains of the cell surface of mouse macrophages, and that this specific lipid environment influences its endocytosis and degradation upon exposure to hepcidin (*i*.*e*. the liver-derived hyposideremic hormone regulating iron metabolism). The authors used MβCD to deplete Chol from cells and consolidate the hypothesis of preferential integration of mouse Fpn1 into lipid rafts (15). Another study revealed that hepcidin binding to human FPN1 is coupled to iron export, with a very important increase in hepcidin affinity in the presence of iron in the secondary iron-binding site of FPN1, which is located within the C-domain in the basal outward-facing conformation (45). The present study confirmed the presence of FPN1 in detergent-resistant fractions of the plasma membrane of human embryonic kidney cells (Figure 1) and unveiled a direct effect of Chol on FPN1 activity (Figure 2). Using MβCD in HEK293 cells stably transfected to overexpress FPN1, we first noticed that Chol depletion significantly reduced FPN1-induced iron export. This negative effect was almost fully reversed after Chol repletion. Unlike Chol, Epi-Chol did not restore iron export, indicating that the effect of Chol is not due to a nonspecific membrane effect, but rather to a stereospecific binding site.

We consequently explored the human FPN1 sequence and structure for possible Chol-binding sites. We used a classical approach, consisting in the identification of CARC, CRAC and CCM motifs. We selected three motifs (out of six) that can be seen on the FPN1 lipid-exposed surface in an outwardly open conformation (which corresponds to the basal state; Figure 3B). In each of these motifs, a central tyrosine was identified as the most important contributor for Chol binding, as the roles played by aromatic side chains in stabilizing the interaction between membrane proteins and Chol are multiple and regarded as critical (30, 46–48), and mutated in either phenylalanine or leucine. Differences were observed between the mutants regarding cell surface expression and iron efflux (Figure 4), but two points are worth discussing. First, the observation of marginal (or even no) effects for substitutions in phenylalanine support the importance of aromatic amino acids for direct interaction with Chol (49). Replacing Tyr29, Tyr220 and Tyr227 with leucine had more pronounced effects on FPN1 activity. Second, the Tyr29Leu mutant, which is predicted to abrogate a CARC motif in the cytoplasmic part of TM1, was found to have a very negative effect on iron egress without altering the plasma membrane expression of FPN1. Importantly, in DRM experiments, the Tyr29Leu, Tyr220Leu and Tyr227Leu mutants shifted to domains of the plasma membrane that are less enriched in Chol. Here again, the Tyr29Leu mutant had the greatest effect (Figure 5).

All these results show that substitutions of central Tyr residues in CARC and CRAC motifs decrease FPN1 localization in Chol-enriched microdomains of the plasma membrane, with a direct consequence on the ability of FPN1 to optimally export iron from cells. The CARC motif in TM1 could play a more important, but not necessarily exclusive, role. In order to obtain further clues, we analyzed the topology of amino acids in the vicinity of Tyr29, Tyr220 and Tyr227 and docked Chol in the corresponding regions. As shown in Figure 6A, Tyr220 and Tyr227 are not positioned colinearly with the aliphatic and basic residues, suggesting a weaker ability for Chol to accommodate with either of the two CRAC motifs in TM6. As a result, docking Chol in this region did not lead to observe poses involving contacts with basic and aromatic side chains (data not shown). On the contrary, the basic, aromatic and aliphatic amino acids of the CARC motif are colinear and located along a groove formed at the interface between helices TM1 and TM5 (Figure 6B). Molecular docking suggests that this groove is well suited to Chol binding. Further computational investigations, especially using molecular dynamics simulations, are needed to assess the stability of Chol in this site and to clarify the atomic details of the Chol-CARC interaction, as well as to elucidate the binding mode of Chol at the level of the two CRAC motifs. We cannot rule out the hypothesis that the binding of Chol on TM6 does not follow the mode adopted by the canonical CRAC/CARC motifs, as already observed (30, 50, 51).

**Figure 6.**
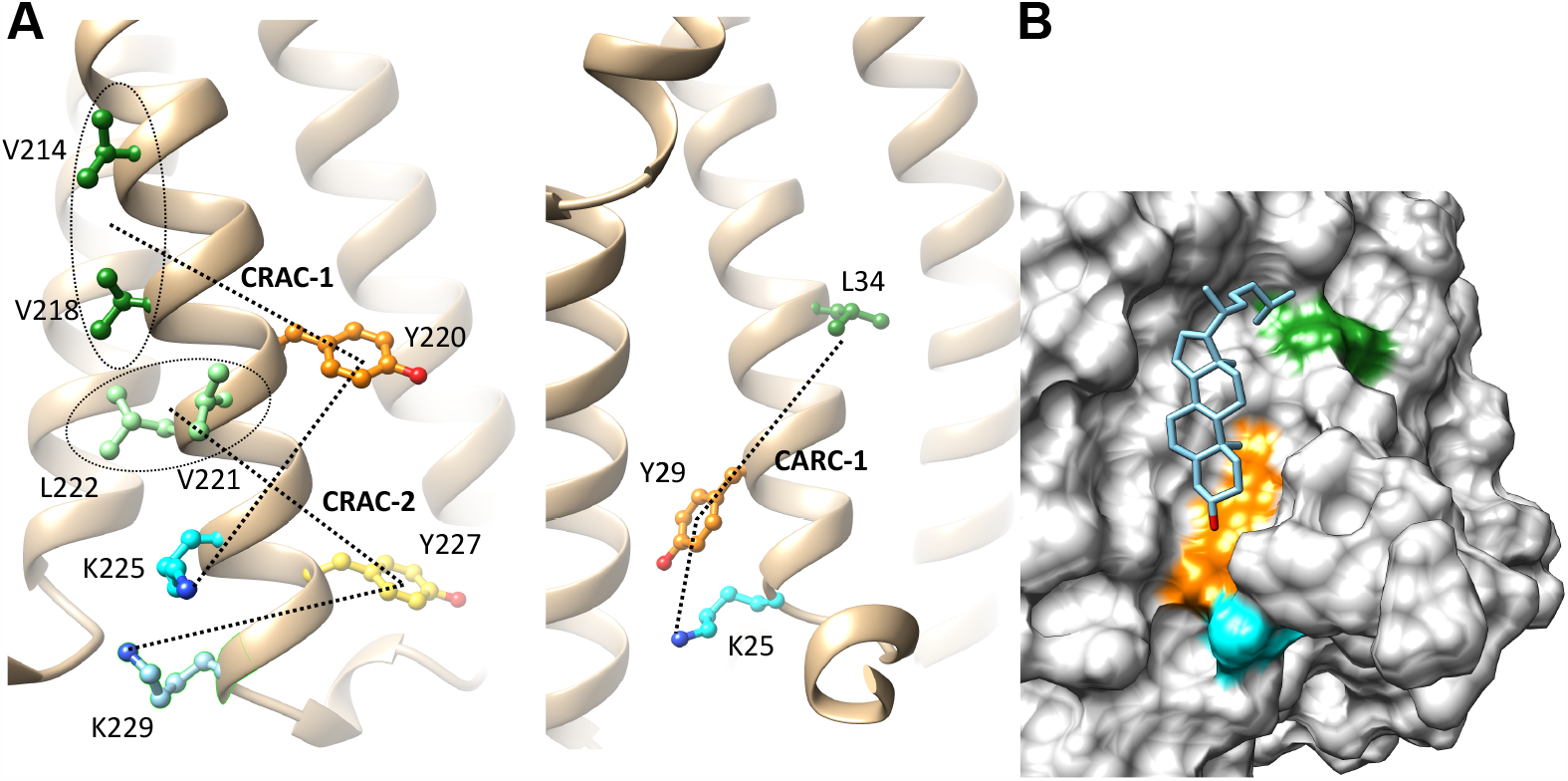
Close-up views of human FPN1 for specific interactions with Chol. (A) Focus on the CARC-1 (right panel) and CRAC-1 and CRAC-2 motifs (left panel), highlighting the positions of their aliphatic, aromatic and basic amino acids. (B) Blind docking of Chol into the CARC-1 motif: surface representation, with the aliphatic, aromatic and basic amino acids shown in green, orange and blue, respectively.

We acknowledge that our current strategy may not be exhaustive, as more Chol-complex protein structures have been studied in recent years, revealing a greater diversity of Chol-binding regions (44, 46, 52). However, we deem relevant to mention that the deep hydrophobic groove identified between TM1 and TM5 has no equivalent in the rest of the protein. This is very similar to what has been reported for another Solute Carrier (SLC) membrane transport protein, namely SLC6A4 or SERT, which is important for reuptake of the neurotransmitter serotine from the synaptic cleft and is known to be modulated by Chol (53).

Multi-pass transmembrane proteins are thought to be less present in ordered membrane domains, which are commonly presented as lipid-rafts (18–22, 54). Some of the available examples indicate that the key determinants of raft affinity for these particular proteins may differ from those of single-pass membrane proteins, which have been more widely studied. Direct interaction with Chol nonetheless remains one of the most decisive factors (54). If we assume that the Tyr29Leu, Tyr220Leu and Tyr226Leu mutations in the TM1-CARC and TM6-CRAC motifs affect association with Chol, the nature of this interaction could influence the plasma membrane lateral organization of FPN1 and its dynamics. Its compartmentalization into domains in which the local concentration of Chol is lower may give rise to rate-limiting conformations, with an impact on iron export. We conversely observed that FPN1 moves towards the most-resistant fractions of the plasma membrane when HEK293T cells are iron loaded (in the same way as earlier studies carried out in mouse macrophages (15)), and therefore require a very efficient FPN1 protein. In this view, Chol may be required as a modulator of FPN1 activity (27, 55). Interestingly, in MFS proteins, TM1 is regarded as a core helix that plays an active role in the formation of thin gates on either side of the lipid bilayer during the clamp-and-switch transport mechanism (18, 19). TM5 is involved in the interface between the N- and C-domains, in tight connection with TM1. We previously revealed that it participates in a network of salt-bridges and hydrogen-bound that is constitutive of the FPN1 intracellular gate and is important for the stability of the iron exporter in its basal outward-facing conformation as well as for transition toward the inward-facing conformation (32). Chol might allosterically influence the conformational dynamics of FPN1, as has been demonstrated for human monoamine SLC6 transporters (not only SERT, but also DAT) in which TM1 and TM5 helices are also considered active contributors of conformational changes (53, 56). Another hypothesis is that FPN1 is packed into Chol- and sphingolipids-enriched vesicles and transported from the Golgi complex to the cell surface, in a way that can promote its arrival into liquid-ordered microdomains (43, 57, 58). A lack of interaction with Chol in the trans-Golgi network could force the inclusion of FPN1 into vesicles with different lipid compositions, further away from lipid-rafts, with a direct consequence on its compartmentalization, but not necessary on its level of expression at the cell surface. Such a situation has been evoked for mutants of the CCM motif of the haemagglutinin of influenza virus (42). These different assumptions, which are not mutually exclusive, will require in depth-investigations.

In summary, we have shown that Chol play an important role in the cell surface localization of human FPN1 in HEK293T and its ability to export iron. Chol is expected to directly bind FPN1 in a way that is likely to promote its integration into liquid-ordered plasma membrane microdomains, and may essentially involve a deep hydrophobic groove at the interface between TM1 and TM5 in the inner leaflet. Sophisticated experimental and computational methods will now be required to provide molecular insights into the interaction between Chol and FPN1.

## Disclosure of Conflicts of Interest

The authors declare that they have no conflicts of interest.

## Acknowledgments

We would like to thank Dr. François Canonne-Hergaux who provided us with the DRM protocol.

## Data availability statement

Data available on request from the authors.

## Notes

**Fundings:** This work was supported by grants from the “Institut Brestois Santé Agro Matière” (IBSAM - AAP2020; Modulofer), the Gaetan Saleun Association and the GR-Ex Laboratory of Excellence (reference ANR-11-LABX-0051). The GR-Ex label is funded by the IdEx “Investissements d’avenir” program of the French National Research Agency (reference ANR-18-IDEX-0001).

### Competing Interest Statement

The authors have declared no competing interest.

