## SupplementaryFigure1 for "Cholesterol modulates the human FPN1 iron export function in plasma membrane liquid-ordered microdomains"

**Supplementary Figure 1.**

A

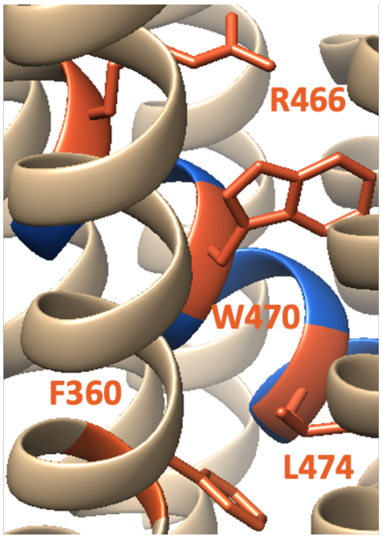


B

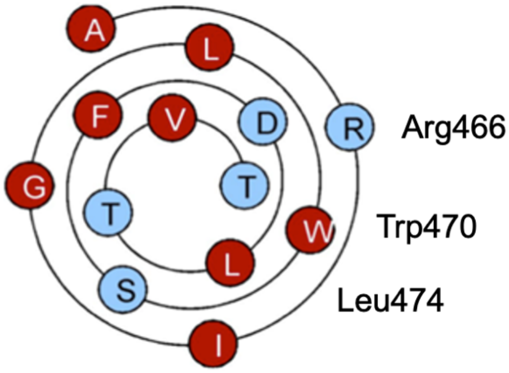


C

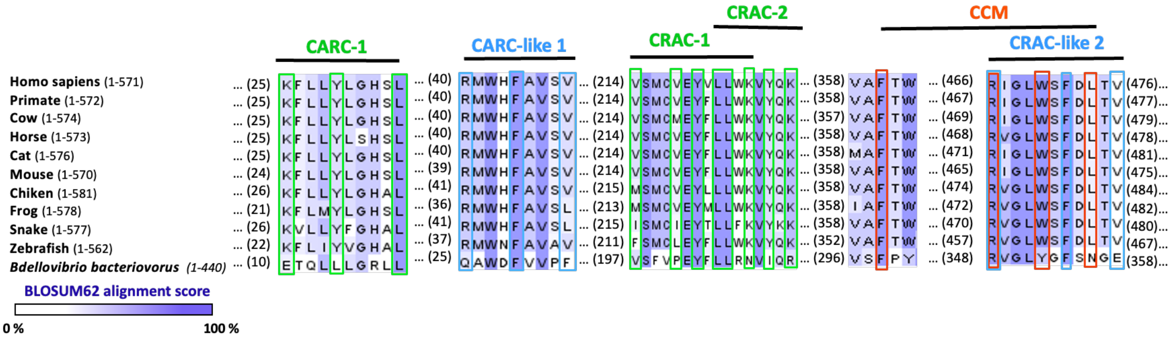

(A) Close-up views of the CCM putative motif within the TM8 and TM10 helices in an outward-facing conformation. Characteristics residues are numbered and colored in orange. (B) Helical wheel plot (Wenxiang diagram; www.*jci-bioinfo.cn/wenxiang2)*, representing the disposition of hydrophobic (red-filled circles) and non-hydrophobic *(*blue-filled circles*) amino acids in TM10.* Position of the characteristic R (Arg466), W (Trp470) and L (Leu474) are specified. (C) Conservation analysis of the putative Chol-binding motifs in different species. The amino acids have been grouped and colored based on the BLOSUM62 matrix. Amino acids forming the CARC/CRAC or CCM motifs are boxed in green and orange, respectively.
